# Membrane remodeling and matrix dispersal intermediates during mammalian acrosomal exocytosis

**DOI:** 10.1101/2021.08.04.455016

**Authors:** Miguel Ricardo Leung, Ravi Teja Ravi, Bart M. Gadella, Tzviya Zeev-Ben-Mordehai

## Abstract

To become fertilization-competent, mammalian sperm must undergo a complex series of biochemical and morphological changes in the female reproductive tract. These changes, collectively called capacitation, culminate in the exocytosis of the acrosome, a large vesicle overlying the nucleus. Acrosomal exocytosis is not an all-or-nothing event, but rather a regulated process in which vesicle cargo disperses gradually. However, the structural mechanisms underlying this controlled release remain undefined. In addition, unlike other exocytotic events, fusing membranes are shed as vesicles; the cell thus loses the entire anterior two-thirds of its plasma membrane and yet remains intact while the remaining non-vesiculated plasma membrane becomes fusogenic. Precisely how cell integrity is maintained throughout this drastic vesiculation process is unclear, as is how it ultimately leads to the acquisition of fusion competence. Here, we use cryo-electron tomography to visualize these processes in unfixed, unstained, fully-hydrated sperm. We show that crystalline structures within the acrosome disassemble during capacitation and acrosomal exocytosis, representing a plausible mechanism for gradual dispersal of the acrosomal matrix. We find that the architecture of the sperm head supports an atypical membrane fission-fusion pathway that maintains cell integrity. Finally, we detail how the acrosome reaction transforms both the micron-scale topography and the nano-scale protein landscape of the sperm surface, thus priming the sperm for fertilization.

**Significance:** Mammalian sperm must undergo a complex series of biochemical and morphological changes in the female reproductive tract in order to become fertilization-competent. These changes culminate in acrosomal exocytosis, during which multiple membrane fusions destabilize the acrosomal vesicle and liberate its contents, which include proteins implicated in penetrating and binding to the egg vestments. Here, we use cryo-electron tomography to visualize acrosomal exocytosis intermediates in unfixed, unstained sperm. Our results suggest structural bases for how gradual dispersal of acrosome contents is regulated, as well as for how the cell remains intact after losing much of its plasma membrane, We also show that acrosomal exocytosis transforms both the micron-scale topography and the nano-scale molecular landscape of the sperm surface, thus priming it for interaction and fusion with the egg. These findings yield important insights into sperm physiology and contribute to our understanding of the fundamental yet enigmatic process of mammalian fertilization.

## Introduction

Mammalian sperm must reside in the female reproductive tract for several hours before they are able to fertilize the egg. During this time, sperm undergo a plethora of biochemical changes collectively called capacitation (Aitken and Nixon, 2013; Bailey, 2010; Gervasi and Visconti, 2016). The discovery of this phenomenon was crucial to the development of in vitro fertilization (Austin, 1951; Chang, 1951, 1959). Capacitation is characterized by cholesterol efflux, phospholipase activation, and altered membrane fluidity, along with a multitude of biochemical changes (Bailey, 2010; Harrison and Gadella, 2005; Nolan and Hammerstedt, 1997; Travis and Kopf, 2002). Together, these changes render sperm capable of undergoing the acrosome reaction, a unique exocytotic event that is an absolute requirement for sperm to become fusion-competent (Yanagimachi, 1981) **(Fig. 1a)**.

**Fig. 1.**
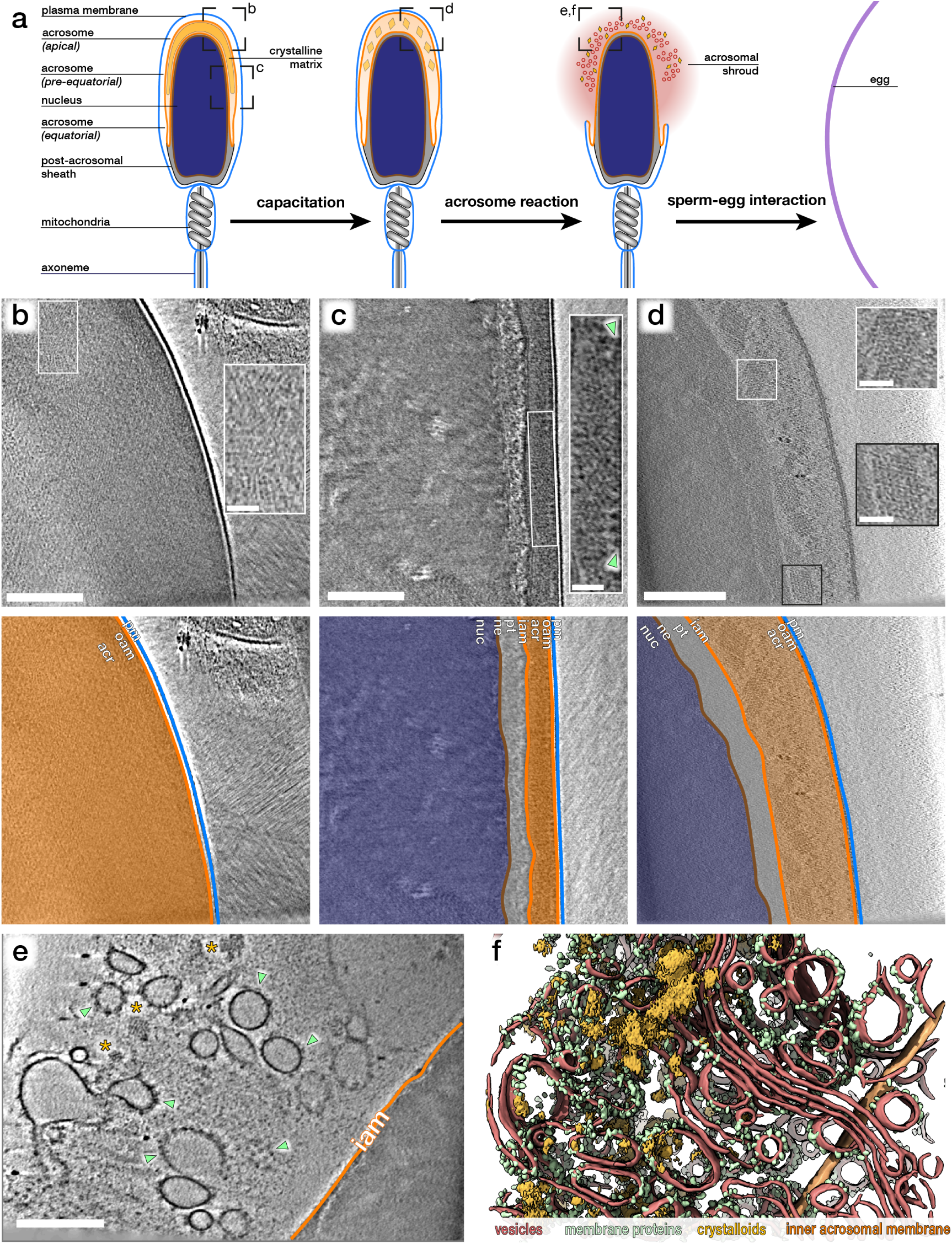
The crystalline fraction of the acrosomal matrix progressively disassembles during acrosomal exocytosis. **(a)** Schematic diagram illustrating the morphological changes that mammalian sperm undergo in the female reproductive tract, simulated in this study in vitro. **(b-c)** Computational slices through Volta phase plate cryo-tomograms of non-capacitated sperm thinned by cryo-focused ion beam milling. Note how the acrosome is dense even in thinned samples. Insets show regions with large crystalloid patches. Large membrane protein densities are visible on the luminal surface of the outer acrosomal membrane (green arrowheads in inset). **(d)** Computational slice through a defocus-contrast cryo-tomogram of a sperm cell whose acrosome had swollen after incubation in capacitating media for ∼2h. Note decondensation of the acrosome and prominent crystalloid patches (insets). **(e-f)** Computational slice (e) and corresponding three-dimensional segmentation (f) of the acrosomal shroud. The cell was incubated in capacitatng media for ∼2h and treated with calcium ionophore for ∼30 min. Note crystalloid patches (asterisks in e, goldenrod in f) and membrane protein-decorated vesicles (green arrowheads in e, green in f). **Scale bars:** 250 nm; insets: 50 nm. **Color scheme:** orange – outer and inner acrosomal membrane, red – vesiculated plasma and outer acrosomal membranes, green – membrane protein densities, goldenrod – crystalloid patches

The acrosome is a large regulated secretory vesicle overlying the anterior two-thirds of the nucleus; its crucial role in mammalian fertilization manifests in the fact that malformation of the acrosome causes infertility in both humans and mice. Three distinct segments of the acrosome can be defined based on their positions along the sperm head **(Fig. 1a)**: the apical segment extends beyond the nucleus and forms the most anterior region of the acrosome; the principal or pre-equatorial segment forms the major part of the acrosome; and the equatorial segment delimits the posterior part of the acrosome. During acrosomal exocytosis, the plasma membrane fuses with the outer acrosomal membrane at multiple points. This destabilizes the acrosome and liberates its contents, which include several proteins implicated in either penetrating through or binding to the egg vestments (Foster and Gerton, 2016).

Acrosomal exocytosis is not an all-or-nothing event but instead involves the gradual dispersal of vesicle cargo (Hardy et al., 1991; Kim and Gerton, 2003; Kim et al., 2001). Biochemical analyses defined two classes of acrosome contents: a soluble fraction that is released shortly after the onset of the acrosome reaction, and a matrix fraction that disperses more slowly. Consistent with this, conventional electron microscopy (EM) studies showed discrete zones within the acrosome, including crystalline material that extends across large areas of the vesicle in sperm from several mammalian species (Fléchon, 2016; Olson and Winfrey, 1994; Phillips, 1972). However, the structural changes in the acrosomal matrix that lead to differential release of acrosome contents are unclear.

Another distinctive feature of acrosomal exocytosis is the lack of membrane recycling. The plasma membrane and the outer acrosomal membrane are shed as vesicles, so the sperm head loses a large portion of its limiting membrane while the remainder becomes fusogenic. Acrosome vesiculation has been studied extensively with classical EM (Barros et al., 1967; Flaherty and Olson, 1991; Jamil and White, 1981; Nagae et al., 1986; Russell et al., 1979; Sosa et al., 2015; Stock and Fraser, 1987; Yudin et al., 1988; Zanetti and Mayorga, 2009), identifying acrosome swelling and membrane docking as clear intermediates leading to exocytosis (Sosa et al., 2015; Tsai et al., 2010; Zanetti and Mayorga, 2009), However, the membrane remodeling pathways that ensure the cell remains intact throughout this dramatic vesiculation process remain undefined.

Here, we use cryo-electron tomography (cryo-ET) to image sperm undergoing in vitro capacitation and acrosome exocytosis. Cryo-ET provides three-dimensional information about rare events in unfixed, unstained, fully-hydrated samples, and thus in close to native conditions (Ng and Gan, 2020). We show that the crystalline component of the acrosomal matrix gradually dissolves during acrosomal exocytosis, explaining the differential release previously observed. We also find that the defined ultrastructural organization of the sperm head facilitates a unique fission-fusion mechanism that maintains cell integrity despite drastic membrane vesiculation. We demonstrate that acrosome exocytosis also facilitates massive membrane protein re-localization onto the post-acrosomal plasma membrane, building a platform for interaction with the egg.

## Results

### In unstimulated sperm, the plasma membrane (PM) and the outer acrosomal membrane (OAM) are closely apposed along the entire acrosome

We plunge-froze sperm from highly fertile, commercial artificial insemination pigs (Sus scrofa domestica) and imaged them using cryo-ET. In intact unstimulated sperm, the plasma membrane (PM) and the outer acrosomal membrane (OAM) are closely apposed along the entire acrosome **(Fig. 1b,c; Fig. S1)**. The PM and the OAM are ∼8-10 nm apart and lie parallel to each other until the equatorial segment of the acrosome, where the vesicle tapers **(Fig. S1d-f)**. This differs from many classical EM images of unstimulated sperm, in which the PM and OAM appear wavy with estimated interbilayer distances of >10 nm (Sosa et al., 2015; Tsai et al., 2010; Zanetti and Mayorga, 2009), and demonstrates the benefits of using cryo-ET to visualise acrosomal exocytosis.

### The crystalline fraction of the acrosomal matrix progressively disassembles during exocytosis

We sought to determine how the internal organization of the acrosome changes during acrosomal exocytosis. We first imaged acro-some contents in intact unstimulated sperm cells thinned to ∼150-200 nm with cryo-focused ion beam (cryo-FIB) milling (Marko et al., 2006; Rigort et al., 2012). Even after thinning and imaging with the Volta phase plate (VPP) (Danev et al., 2014; Fukuda et al., 2015), the acrosome lumen was still very dense **(Fig. 1b,c)**. Nonetheless, we observed large patches of crystalline material in both the apical and pre-equatorial regions of the acrosome **(insets in Fig. 1b,c)**, similar to the structures observed in rat sperm (Phillips, 1972), rabbit sperm (Olson and Winfrey, 1994), and ram sperm (Fléchon, 2016).

To visualise acrosomal exocytosis, we imaged sperm incubated in capacitating media (containing calcium, bicarbonate, and bovine serum albumin) for ∼2 h and subsequently treated with calcium ionophore A23187 **(Fig. S2)** or progesterone for up to 30 min. Following treatment, fully acrosome-reacted sperm could be readily targeted in low-magnification cryo-EM projection images **(Fig. S3)**. These cells were surrounded by a cloud of vesicles (the acrosomal shroud) **(Fig. S3a)** and their apical regions had become very thin due to the loss of the acrosome **(Fig. S3b)**, allowing us to image them without cryo-FIB milling. We note that acrosomal shrouds tend to remain associated with sperm heads despite several pipetting and dilution steps before imaging. In our analyses, we excluded sperm in which the plasma membrane peeled off at the equatorial/post-acrosomal regions, which would likely result in loss of cell integrity and thus in cell death **(Fig. S3c-d)**.

We found that the acrosomal shroud consists of a highly heterogeneous population of vesicles that are decorated with membrane proteins **(Fig. 1e,f)**. Interspersed between these vesicles are the contents of the acrosome, including striking crystalloid patches that were heterogeneous in both size and shape **(Fig. 1e; Fig. S4a,b)** (ionophore: 12/13 tomograms, each from a different cell, from 3 different animals; progesterone: 5/5 tomograms, each from a different cell, from 1 animal).

To follow the crystalloids during intermediate stages of exocytosis, we imaged cells incubated in capacitating media without ionophore treatment. Crystalloids were readily visible in swollen acrosomes **(Fig. 1d)**, where they had already begun to dissociate into smaller fragments. We then used subtomogram averaging to resolve the structure of the crystalloid patches at a resolution of ∼30-40 Å **(Fig. 2)**. We chose to average from capacitated sperm since the surrounding material was too dense in naïve sperm. The patches were also larger in capacitated sperm than in the shrouds of acrosome-reacted sperm, which facilitated averaging by increasing particle numbers. Our averages reveal that the crystalloids adopt a tetragonal body-centered crystal lattice with apparent unit cell dimensions a = 28 nm, b = 28 nm, c = 40 nm.

**Fig. 2.**
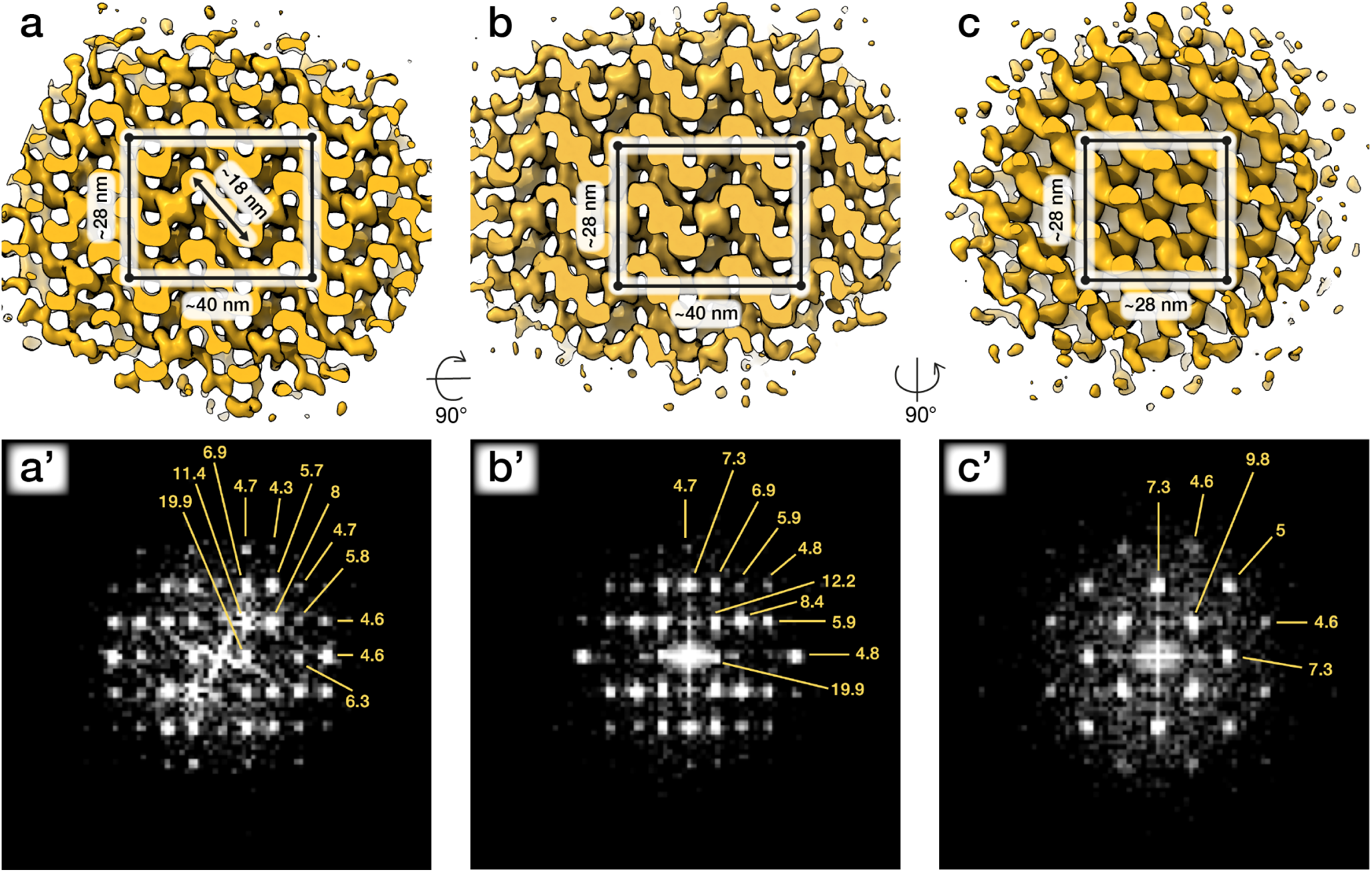
*In situ* structures of crystalloid patches from capacitated sperm. Three orthogonal isosurface views **(a-c)** and corresponding Fourier transforms **(a’-c’)** of a subtomogram average of 40 × 40 × 40 nm crystalloid patch from swollen acrosomes of boar sperm incubated in capacitating media for ∼2 h. The putative tetragonal body-centered unit cell is annotated in (a-c). Averages were generated from ∼1400 particles from two cells.

Taken together, our data indicate that the crystalloids in the acrosomal shroud result from disassembly of an initial larger superstructure. These crystalloids may thus represent the core of the acrosomal matrix, acting as a structural scaffold onto which soluble components of the acrosome are anchored. Their progressive disassembly may represent a mechanism for controlled release of acrosome contents that appears to be conserved across mammals.

### An atypical membrane fission-fusion pathway maintains cell integrity at the equatorial segment

We then sought to trace membrane remodelling intermediates involved in capacitation and acrosomal exocytosis. Sperm within an ejaculate are inherently variable (Buffone et al., 2014), which precludes a strictly timepoint-based assessment of the reaction coordinate. We instead imaged several cells from several different animals **(Table S1)**, analysed the dataset for membrane remodelling intermediates that we could observe consistently, and ordered these stages relative to one another **(Fig. 3, Fig. 4)**.

**Fig. 3.**
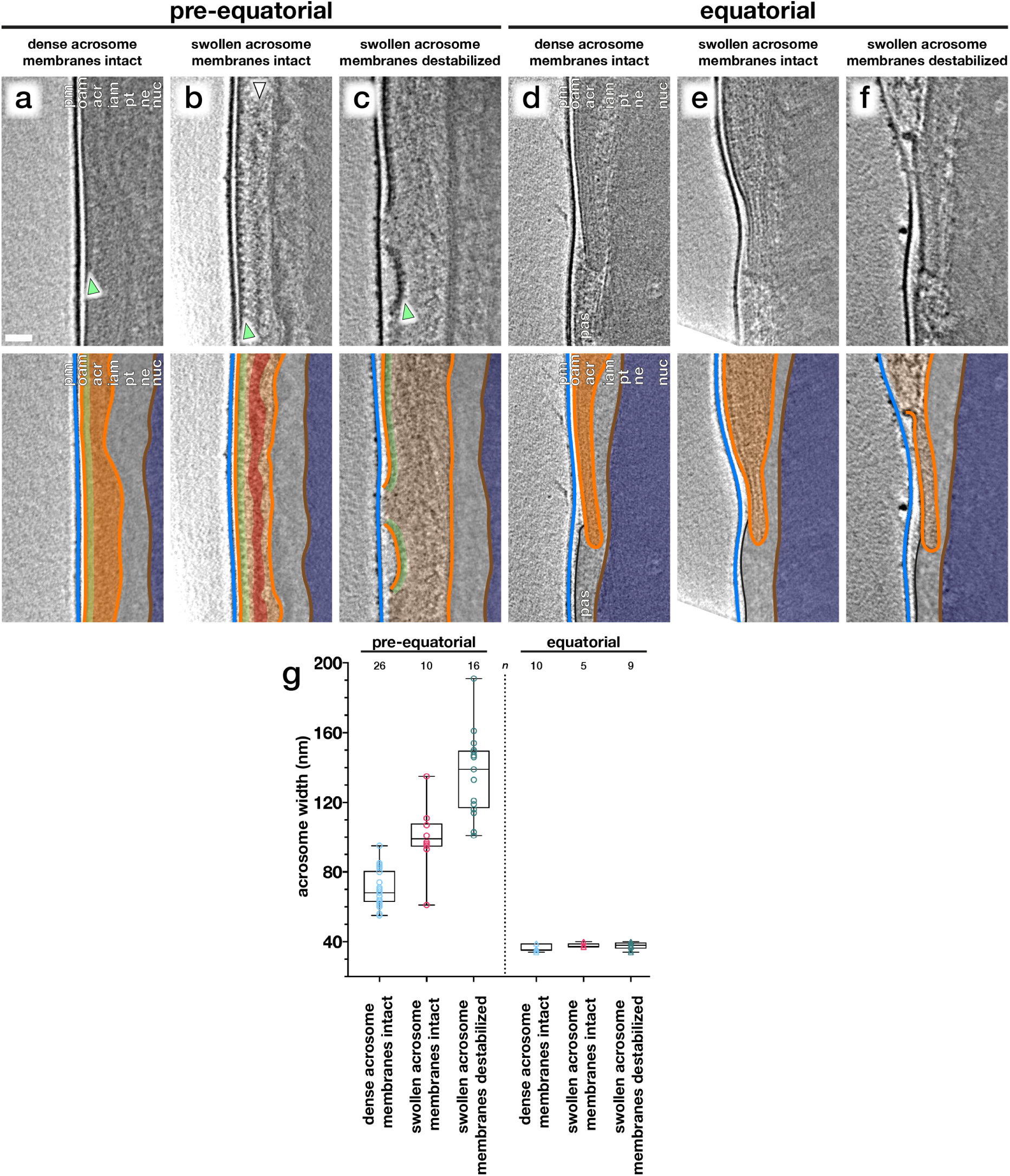
Acrosome swelling is associated with membrane destabilization in capacitated sperm. **(a-f)** Computational slices (top panels) and schematic annotations (bottom panels) of Volta phase plate cryo-tomograms of the pre-equatorial (a-c) and equatorial (d-f) region of sperm before (a,d) or after (b-c, e-f) a ∼2 h incubation in capacitating media without ionophore stimulation. Note the rows of membrane protein densities on the outer acrosomal membrane (green arrowheads in a-c) and the streak of acrosomal matrix that remains condensed during acrosome swelling (white arrowhead in b). **(g)** Measurements of acrosomal width at the pre-equatorial (left) and equatorial (right) regions in sperm showing various states of acrosome swelling and membrane destabilization. For each tomogram, acrosome width was measured as the distance between the outer and inner acrosomal membranes at three different locations. Each tomogram is from a different cell (n indicates the number of tomograms used for analysis). The boxes indicate the median and interquartile range, while the whiskers indicate the minimum and maximum values. **Scale bar:** 50 nm. **Labels:** pm – plasma membrane, oam – outer acrosomal membrane, acr – acrosome, iam – inner acrosomal membrane, pt – perinuclear theca, ne – nuclear envelope, nuc – nucleus, pas – post-acrosomal sheath. **Color scheme:** blue – plasma membrane, orange – outer and inner acrosomal membranes, green – membrane protein densities, red – condensed acrosomal material, light grey – perinuclear theca and post-acrosomal sheath, brown – nuclear envelope, dark blue – nucleus

**Fig. 4.**
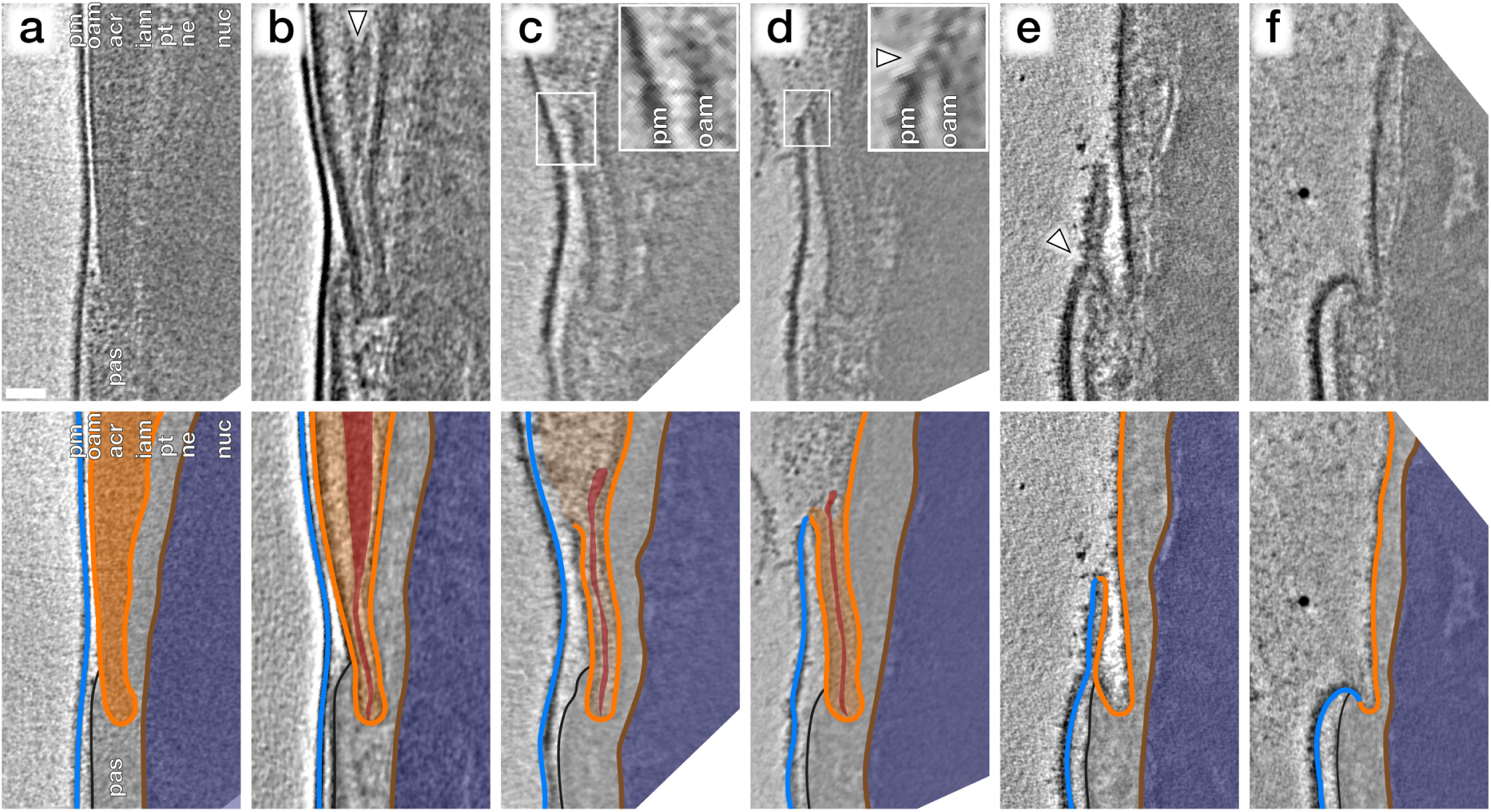
An atypical membrane fission-fusion pathway maintains cell integrity at the equatorial region. **(a-f)** Computational slices (top panels) and corresponding schematic annotations (bottom panels) of Volta phase plate cryo-tomograms of the equatorial region of boar sperm incubated for ∼2 h in capacitating media without (a-d) and with (e-f) subsequent ∼30 min ionophore stimulation. In (b), the white arrowhead indicates condensed material at the core of the acrosome that is continuous with a thin streak of electron-dense material in the equatorial segment. In (c), the inset shows the point of OAM rupture immediately anterior to the equatorial region. In (d), the inset shows that the OAM has fused (white arrowhead) with the overlying PM at this location. **Scale bar:** 50 nm. **Labels:** pm – plasma membrane, oam – outer acrosomal membrane, acr – acrosome, iam – inner acrosomal membrane, pt – perinuclear theca, ne – nuclear envelope, nuc – nucleus, pas – post-acrosomal sheath’ **Color scheme:** blue – plasma membrane, orange – outer and inner acrosomal membranes, light grey – perinuclear theca and post-acrosomal sheath, brown – nuclear envelope, dark blue – nucleus

We captured a range of intermediates already in capacitated sperm without ionophore stimulation **(Fig. 3)**, including acrosome swelling, membrane docking, and membrane destabilization **(Fig. S5)**. Acrosome swelling is one of the earliest stages of acrosomal exocytosis (Sosa et al., 2015; Zanetti and Mayorga, 2009), and indeed was observed even in capacitated cells (Boerke et al., 2014). Swelling is associated with decondensation of acrosomal contents, but our tomograms reveal that decondensation is not uniform. Specifically, a dense core remains near the center of the vesicle **(white arrowhead in Fig. 3b)**, which is continuous with a thin streak of electron-dense material sandwiched between the OAM and the IAM at the end of the acrosome **(white arrowhead in Fig. 4b)**. Decondensation also improves contrast in the acrosome, making visible large membrane protein densities on the luminal surface of the OAM **(green arrowhead in Fig. 3b**,**c)**. These structures are also visible in VPP tomograms of FIB-milled unstimulated sperm **(green arrowheads in Fig. 1b inset and Fig. 3a)**. The OAM proteins form rows of teeth-like densities, each extending ∼14 nm into the OAM lumen and spaced ∼18 nm from its neighbours.

We could further distinguish between two stages of acrosome swelling: one in which the acrosome swells but the overlying membranes are still intact **(Fig. 3b,e)**, and another in which the OAM already destabilizes **(Fig. 3c,f)**. OAM destabilization is characterised by local membrane rupture **(Fig. 3c,f; Fig. 4c; Fig. S5g-l)**. OAM rupture was associated with the extent of acrosome swelling; in cells with ruptured OAMs, the acrosome had swollen to nearly twice its original width (136 ± 24 nm versus 71 ± 11 nm, with an intermediate value of 100 ± 18 nm in cells with swollen acrosomes but intact membranes) **(Fig. 3g)**. In contrast, the width of the equatorial segment did not change significantly even in cells with ruptured OAMs (acrosome swollen, membranes destabilized: 38 ± 2 nm; acrosome swollen, membranes intact: 38 ± 1 nm; dense acrosome, membranes intact: 36 ± 2 nm). Notably, OAM rupture occurred just anterior to the equatorial segment, on average 260 ± 80 nm from the end of the acrosome (mean ± s.d., 8 tomograms, each from a different cell) **(Fig. 3f)**.

Focusing on the equatorial segment, we observed an atypical membrane fission-fusion pathway that mediates resealing of the sperm head **(Fig. 4)**. The ruptured end of the OAM fuses with the overlying PM **(Fig. 4c,d; Fig. S5j-l)** and after this fusion event, the electron-dense streak also diffuses, leaving a hairpin-shaped membrane **(Fig. 4e, Fig. S6a-b)**. This hairpin-shaped membrane then constricts and buds off just anterior to the post-acrosomal sheath **(Fig. 4e, arrowhead)**, yielding the characteristic morphology of acrosome-reacted cells **(Fig. 4f, Fig. S6c)**.

### Acrosomal exocytosis transforms the molecular landscape of the sperm plasma membrane

After loss of the acrosome, the inner acrosomal membrane (IAM) is the new limiting membrane of the apical segment of the sperm cell **(Fig. 1e)**. The PM overlying the equatorial/post-acrosomal segment remains intact and is now continuous with the IAM **(Fig. 5)**. Segmentation of high-contrast tomograms acquired with a VPP revealed that the PM forms a “sheath” around the post-acrosomal region **(Fig. 5d, Fig. S6)**. Our tomograms also showed tubulovesicular projections overlying the equatorial segment **(Fig. 5d, Fig. S6)**. These tubular membranes are consistent with those observed by freeze-fracture EM (Aguas and da Silva, 1989). Thus, the acrosome reaction remodels the overall topography of the sperm surface.

**Fig. 5.**
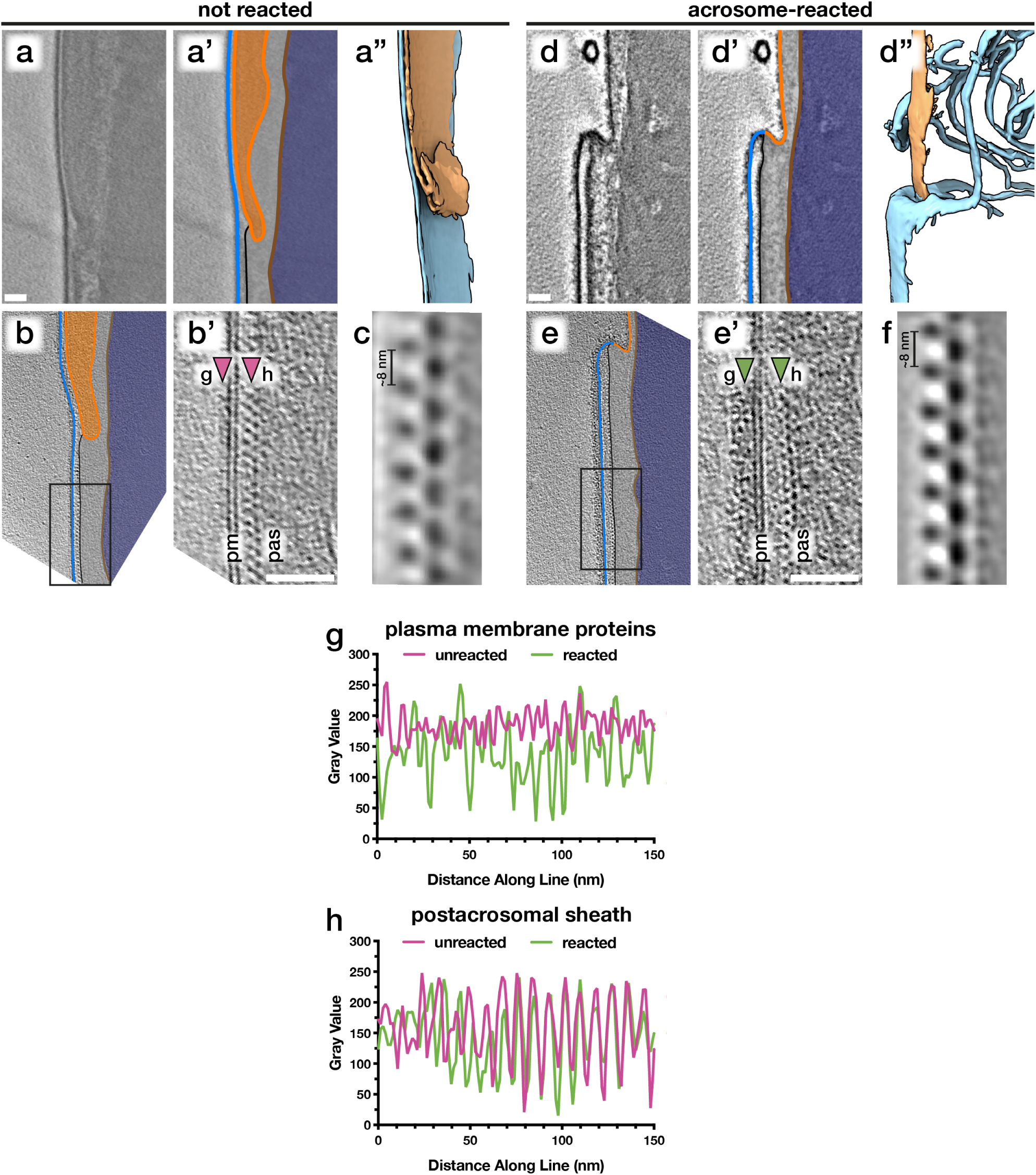
Acrosomal exocytosis transforms the molecular landscape of the sperm plasma membrane. Comparing the post-acrosomal plasma membrane in unreacted **(a-c)** versus acrosome-reacted **(d-f)** reveals major differences in membrane protein decoration. **(a**,**d)** Computational slices (a & a’, d & d’) and corresponding segmentations (a”, d”) of VPP cryo-tomograms illustrating how acrosomal exocytosis remodels the topography of the equatorial region. **(b**,**e)** Computational slices through defocus-contrast cryotomograms illustrating how the post-acrosomal plasma membrane becomes densely-packed with membrane proteins after acrosomal exocytosis. **(c**,**f)** Subtomogram averages of the post-acrosomal sheath from unreacted (c) and acrosome-reacted (f) cells. **(g**,**h)** Exemplary linescans illustrating changes in membrane protein density after the acrosome reaction (g), as well as the lack of noticeable change in the post-acrosomal sheath (h). Linescans were taken at the approximate locations marked by arrowheads in (b’) and (e’). **Scale bars:** 50 nm. **Labels:** pm – plasma membrane, pas – post-acrosomal sheath. **Color scheme:** blue – plasma membrane, orange – outer and inner acrosomal membranes, light grey – perinuclear theca and post-acrosomal sheath, brown – nuclear envelope, dark blue – nucleus

We then compared the post-acrosomal segment in naïve versus acrosome-reacted sperm **(Fig. 5b,e; Fig. S7)**. Inspecting the PM overlying the post-acrosomal sheath reveals major differences in protein decoration. In unreacted cells, the post-acrosomal PM is relatively smooth, with only a few small protein densities protruding from the membrane **(Fig. 5b, Fig. S7a-c)** (14/14 tomograms, each from a different cell, from 6 different animals). In contrast, the post-acrosomal PM was densely packed with membrane protein densities in ∼80% of tomograms of acrosome-reacted cells (30/37 tomograms, each from a different cell, from 5 different animals) **(Fig. 5e,g; Fig. S7d-f)**. These densities do not appear to be ordered, which suggests that they represent a range of different conformations, proteins, or protein complexes. Thus, the acrosome reaction results in massive membrane protein relocalization that alters the molecular landscape of the sperm surface.

We then used subtomogram averaging to define the sub-structure of the post-acrosomal sheath in more detail **(Fig. 5c,f)**. The peripheral layer of the post-acrosomal sheath, immediately underlying the PM, consists of a multi-layered structure with an ∼8-nm repeating unit. Our averages reveal that neither the substructure nor the overall organization of the post-acrosomal sheath change noticeably after acrosomal exocytosis, which is consistent with measurements directly from tomograms **(Fig. 5h)**. Similarly, the distance between the PM and the post-acrosomal sheath remains relatively unchanged (unreacted: 15 ± 2 nm, reacted: 18 ± 3 nm).

## Discussion

### Gradual disassembly of the crystalline matrix may represent a mechanism for controlled release of acrosome contents

Leading up to and during acrosomal exocytosis, acrosome contents disperse at rates dependent on their partitioning into either soluble or particulate fractions. The particulate fraction, also known as the acrosomal matrix, disperses gradually in a process dependent on alkalization and proteolytic self-digestion (Buffone et al., 2008). However, we do not understand the underlying structural transitions in the acrosomal matrix that regulate the dispersal of acrosomal contents. Studies of acrosomal matrix dispersal are often performed on guinea pig sperm, which have large acrosomes partitioned into subdomains easily visible by transmission EM (Flaherty and Olson, 1988; Hardy et al., 1991; Kim et al., 2001; Olson and Winfrey, 1994). Here, we use cryo-ET to show that the acrosome is structurally compartmentalized also in boar sperm **(Fig. 1)**, which have comparatively thin acrosomes and no obvious subdomains when viewed by conventional EM (Buffone et al., 2008).

Specifically, we find an extensive crystalline fraction in the boar sperm acrosome. Although crystalline structures have been demonstrated previously in acrosomes of other mammals, they were not followed throughout capacitation and acrosomal exocytosis. Our data now show that the crystalline fraction begins to disassemble during capacitation **(Fig. 1d)** and continues to do so during acrosomal exocytosis, resulting in small patches scattered amongst the vesicles of the acrosomal shroud **(Fig. 1e,f)**. Gradual disassembly of the crystalline matrix thus represents a plausible mechanism for controlled release of acrosome contents. Refining this model will require further studies aimed at determining the nature of the crystalline fraction – for instance, whether it represents a scaffolding structure or a storage phase for inactive enzymes. The subtomogram averages we present here **(Fig. 2)** can help towards this goal by providing constraints on the molecular dimensions of candidate proteins.

### Cryo-ET reveals how capacitation-associated membrane destabilization relates to the membrane fission-fusion processes involved in acrosomal exocytosis

Our observations suggest that acrosome swelling relates directly to membrane destabilization **(Fig. 3)**, which is likely caused by an increase in membrane tension in addition to known changes in membrane composition mediated by cholesterol efflux and phospholipase activation (Aitken and Nixon, 2013; Asano et al., 2013). A role for swelling-dependent membrane destabilization is also supported by observations that hyper-osmotic conditions inhibit the acrosome reaction (Bielfeld et al., 1993) and that lysophosphatidylcholine, a positive curvature amphiphile that reduces the energetic barrier for membrane rupture (Glushakova et al., 2005), promotes the reaction (de Lamirande et al., 1997; Parrish et al., 1988). Meanwhile, the equatorial region is stabilized by the electron-dense core of the acrosome **(Fig. 3d-f)** and the post-acrosomal region likewise stabilized by the post-acrosomal sheath. Thus, the precise organization of the sperm head facilitates the rupture-fusion pathway that maintains cell integrity despite destabilization and vesiculation of the rest of the acrosome **(Fig. 4)**.

The intermediates we observe do not appear to fit the canonical fusion-by-hemifusion pathway; instead, the presence of membrane edges is reminiscent of the rupture-insertion pathway (Chlanda et al., 2016; Haldar et al., 2019). Whether fusion proceeds via hemifusion or via rupture-insertion depends on membrane spontaneous curvature and hence on lipid composition, with the rupture-insertion pathway strongly favoring cholesterol-poor bilayers (Chlanda et al., 2016; Haldar et al., 2019). This may be particularly relevant given that one of the molecular signatures of capacitation is cholesterol efflux.

### Acrosomal exocytosis transforms the molecular landscape of the sperm plasma membrane

Acrosomal exocytosis is an absolute requirement for mammalian sperm to fuse with the egg (Yanagimachi, 1981). Fluorescence microscopy has shown that as a result of the acrosome reaction, Izumo1 re-localizes onto the plasma membrane, allowing it to interact with its oocyte-borne partner, Juno, to mediate sperm-egg adhesion (Satouh et al., 2012). However, the Izumo1-Juno interaction is not sufficient to mediate membrane fusion (Bianchi et al., 2014). Our understanding of sperm-egg fusion is hampered by the fact that, beyond the translocation of Izumo1, we know very little about what happens to the molecular landscape of the sperm surface after the acrosome reaction.

Here, we show that acrosomal exocytosis transforms both the micron-scale topography and the molecular landscape of the sperm surface **(Fig. 5)**. We find that the post-acrosomal plasma membrane becomes heavily decorated with membrane protein densities. Such changes may be due to the re-localization of membrane proteins, similar to the phenomenon observed by freeze-fracture EM for the acrosomal cap region (Aguas and da Silva, 1989), or to the binding of liberated acrosomal proteins to pre-existing receptors. The post-acrosomal membrane protein densities do not appear to be ordered, which suggests that they represent a range of different conformations, proteins, or protein complexes. Indeed, in addition to Izumo1, there are now a number of proteins on mammalian sperm that are known to be essential for sperm-egg binding and fusion (Fujihara et al., 2020; Inoue et al., 2005; Lamas-Toranzo et al., 2020; Noda et al., 2020). Our results therefore complement the emerging view that mammalian sperm-egg fusion involves several molecular species acting in concert. Our work also opens avenues for future work into how these various players are organized on the sperm membrane at the nano-scale.

## Acknowledgements

The authors thank Dr. M Vanevic for excellent computational support, and Dr. SC Howes, Ingr. CTWM Schneijdenberg & JD Meeldijk for managing and maintaining the Utrecht University EM Square facility. The authors also thank S Leemans and L Teeuwen for their help with sperm preparation in initial stages of the project. The authors also thank the Henriques Lab for the publicly-available LATEX template. This work benefitted from access to the Netherlands Center for Electron Nanoscopy (NeCEN) with support from operators Dr. RS Dillard and Dr. CA Diebolder and IT support from B Alewijnse. This work was funded by NWO Start-Up Grant 740.018.007 to TZ, and MRL is supported by a Clarendon Fund-Nuffield Department of Medicine Prize Studentship.

## Author Contributions

MRL and RTR prepared samples for cryo-EM. MRL, RTR, and TZ collected and analyzed cryo-ET data. MRL, RTR, BMG, and TZ wrote the manuscript.

## Materials and Methods

### Sperm washing, capacitation, and acrosome reaction

Freshly-ejaculated pig (Sus scrofa domestica) semen was purchased from an artificial insemination company (AIM Varkens KI Nederland). Semen was typically diluted in Beltsville’s thawing solution (BTS: 205 mM glucose, 20 mM NaCl, 5 mM KCl, 15 mM NaHCO3, 3 mM EDTA) and stored at 18 °C until use. Sperm were used within 1 day of delivery. Sperm were gently layered onto a discontinuous gradient consisting of 2 mL of 70% Percoll overlaid with 4 mL of 30% Percoll, both in HEPES-buffered saline (HBS: 20 mM HEPES, 137 mM NaCl, 10 mM glucose, 2.5 mM KCl, 0.1% kanamycin, pH 7.6) and centrifuged at 750 g for 15 min. Pelleted cells were washed once in phosphate-buffered saline (PBS: 137 mM NaCl, 3 mM KCl, 8 mM Na2HPO4, 1.5 mM KH2PO4, pH 7.4), resuspended in PBS, and counted.

Washed sperm were resuspended in 1X TALP (20 mM HEPES, 90 mM NaCl, 21.7 mM sodium lactate, 15 mM NaHCO3, 5 mM glucose, 3.1 mM KCl, 2 mM CaCl2, 1 mM sodium pyruvate, 0.4 mM MgSO4, mM NaH2PO4, 100 µg/mL kanamycin, 0.3% w/v fatty acid-free BSA (Sigma), pH 7.4) at ∼10-20 × 106 cells/mL. Sperm were allowed to capacitate for between 2 and 2.5 hours at 37 °C, 5% CO2. In order to stimulate the acrosome reaction more rapidly and in a larger percentage of cells, either calcium ionophore A23187 (Sigma) or progesterone (Sigma) was added to capacitated cells to a final concentration of either 5 µM or 3 µM, respectively. Cells were incubated for a further 30 min (for ionophore) or 1 h (for progesterone) at 37 °C, 5% CO2.

For flow cytometry, cells were first washed with PBS and their concentration adjusted to 30-50 × 106 cells/mL. Sperm were then stained with propidium iodide (PI) (LifeTechnologies) and with PNA-fluorescein isothiocyanate (FITC)(Sigma), both at a final concentration of 1 µg/mL. Sperm were then diluted 1/100 to 0.5 × 106 cells/mL and analyzed using a BD FACSCanto II flow cytometer. Viable, acrosome-reacted cells were defined as those in the PI+FITC+ quadrant of the cytogram.

### Cryo-EM grid preparation

Typically, 3 µL of a suspension containing either 2-3 × 106 cells/mL (for whole cell tomography) or 20-30 × 106 cells/mL (for cryo-FIB milling) was pipetted onto a glow-discharged Quantifoil R 2/1 200- mesh holey carbon grid. One µL of a suspension of BSA-conjugated gold beads (Aurion) was added, and the grids then blotted manually from the back (opposite the side of cell deposition) for ∼3 s (for whole cell tomography) or for ∼5-6 s (for cryo-FIB milling) using a manual plunge-freezer (MPI Martinsreid). Grids were immediately plunged into a liquid ethane-propane mix (37% ethane) (Tivol et al., 2008) cooled to liquid nitrogen temperature. Grids were stored under liquid nitrogen until imaging.

### Cryo-focused ion beam milling

Grids were mounted into modified Autogrids (ThermoFisher) for mechanical support. Clipped grids were loaded into an Aquilos (ThermoFisher) dual-beam cryo-focused ion beam/scanning electron microscope (cryo-FIB/SEM). All SEM imaging was performed at 2 kV and 13 pA, whereas FIB imaging for targeting was performed at 30 kV and 10 pA. Milling was typically performed with a stage tilt of 18°, so lamellae were inclined 11° relative to the grid. Each lamella was milled in four stages: an initial rough mill at 1 nA beam current, an intermediate mill at 300 pA, a fine mill at 100 pA, and a polishing step at 30 pA. Lamellae were milled with the wedge pre-milling technique described in (Schaffer et al., 2017) and with expansion segments as described in (Wolff et al., 2019).

### Tilt series acquisition

Tilt series were acquired on either a Talos Arctica (ThermoFisher) operating at 200 kV or a Titan Krios (ThermoFisher) operating at 300 kV, both equipped with a post-column energy filter (Gatan) in zero-loss imaging mode with a 20-eV energy-selecting slit. All images were recorded on a K2 Summit direct electron detector (Gatan) in either counting or super-resolution mode with dose-fractionation. Tilt series were collected using SerialEM (Mastronarde, 2005) at a target defocus of between -4 and -6 µm (conventional defocus-contrast) or between -0.5 and -1.5 µm (for tilt series acquired with the Volta phase plate). Tilt series were typically recorded using either strict or grouped dose-symmetric schemes, either spanning ± 56° in 2° increments or ± 54° in 3° increments, with total dose limited to ∼100 e-/Å2.

### Tomogram reconstruction

Frames were aligned either post-acquisition using Motioncor2 1.2.1 (Zheng et al., 2017) or on-the-fly using Warp (Tegunov and Cramer, 2019). Frames were usually collected in counting mode; when super-resolution frames were used, they were binned 2X during motion correction. Tomograms were reconstructed in IMOD (Kremer et al., 1996) using weighted back-projection, with a SIRT-like filter (Zeng, 2012) applied for visualization and segmentation. Defocus-contrast tomograms were CTF-corrected in IMOD using ctfphaseflip while VPP tomograms were left uncorrected.

### Tomogram segmentation

Tomogram segmentation was generally performed semi-automatically. Initial segmentation was performed using the neural network-based TomoSeg package in EMAN 2.21. Segmentation was then refined manually in either Avizo 9.2.0 (FEI) or Chimera 1.12. Membrane distance measurements were performed using built-in functions in Avizo 9.2.0.

### Subtomogram averaging of crystalloid patches

Subtomogram averaging with missing wedge compensation was performed using PEET 1.13.0 (Heumann et al., 2011; Nicastro et al., 2006). Alignments were performed first on 4x-binned defocus-contrast tomograms, after which aligned positions and orientations were transferred to 2x-binned data using scripts shared by Dr. Daven Vasishtan.

Particle positions were seeded by generating a three-dimensional grid of points in crystalloid patches using the gridInit program. All particle orientations were randomized and initial alignments allowed for full rotational searches around all axes. To ensure consistency, two independent initial alignments were performed, each using a randomly-selected particle from a separate tomogram as an initial reference. Since alignments converged on a similar structure, alignments were continued. The dataset was cleaned by removing all particles with a cross-correlation value less than one standard deviation above the mean (which removed poorly-aligning particles such as those at the edges of crystalloid patches) and by (ii) removing overlapping particles. The orientations of the remaining particles were again randomized and another alignment performed. After a final particle clean-up by classification, a final restricted alignment run was performed. Averages presented in the manuscript were filtered to the estimated resolution based on the Fourier shell correlation (FSC) at a cut-off of 0.5 (Nicastro et al., 2006).

### Measurements and quantification

All measurements were performed on ∼20-nm thick central tomographic slices. Acrosomal width was measured manually in IMOD as the distance between the outer and inner acrosomal membranes. For each tomogram, three measurements were recorded at different locations to account for slight variations in the shape of the acrosome. Linescans for measurement of the post-acrosomal sheath were performed in Fiji v 2.0.0-rc-69/1.52p.

## Data availability

Subtomogram average maps will be deposited to the Electron Microscopy Data Bank (EMDB).

---

**Fig. S1.**
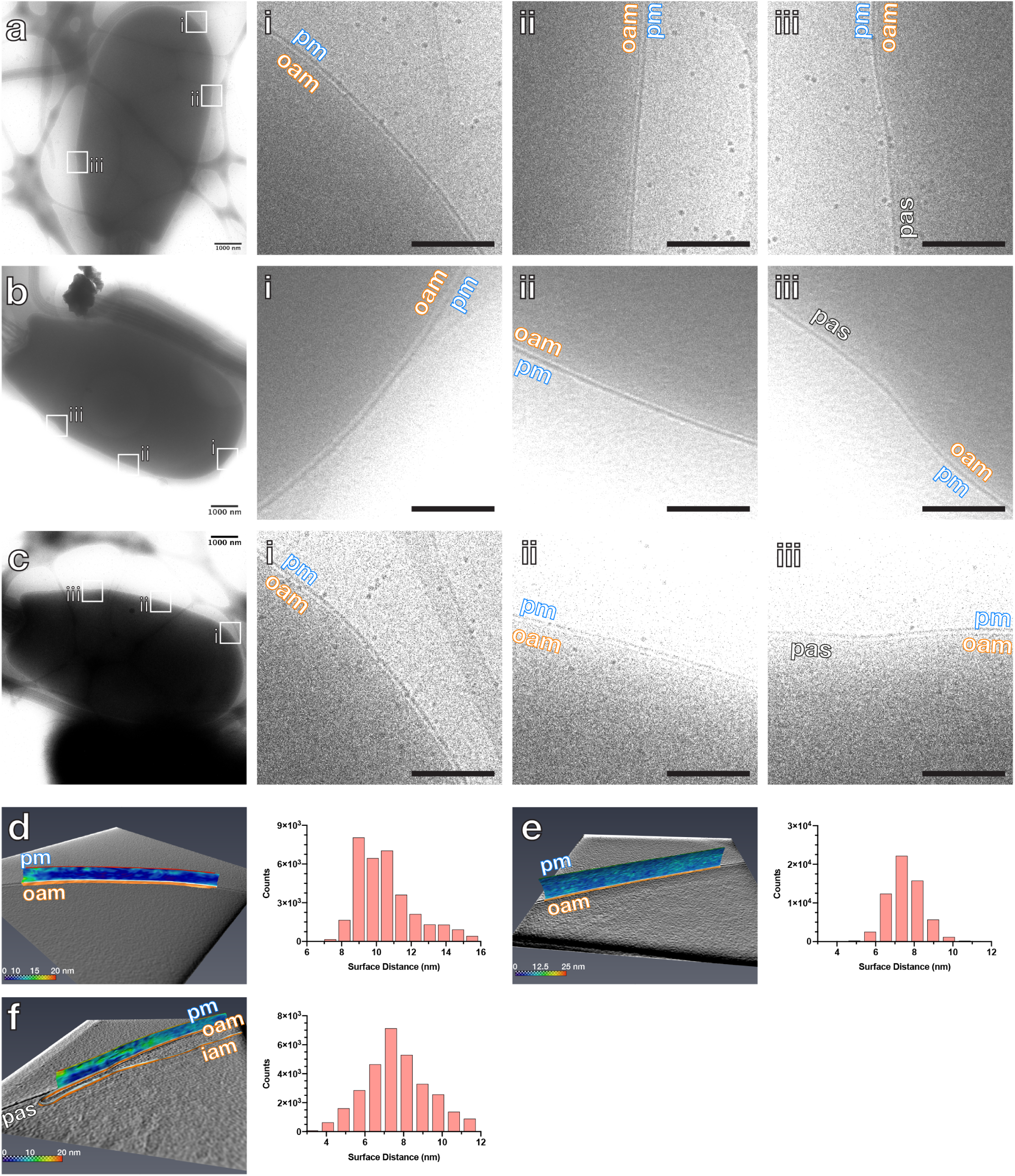
The plasma membrane and the outer acrosomal membrane are closely spaced even in non-capacitated sperm. **(a-c)** Low-magnification high-dose cryo-EM projection images of non-capacitated boar sperm heads. Digital zooms show close apposition between the plasma membrane and the outer acrosomal membrane at the apical (i), pre-equatorial (ii), and equatorial (iii) regions. **(d-f)** Left panels show three-dimensional segmentations of the plasma membrane and the outer acrosomal membrane at the apical (d), pre-equatorial (e), and equatorial (f) regions. The plasma membrane is colored based on distance from the outer acrosomal membrane, which is shown as a cutaway view in orange. Right panels show corresponding histograms of intermembrane distances. **Scale bars:** (a-c) 1 µm; digital zooms: 250 nm. **Labels:** pm - plasma membrane, oam - outer acrosomal membrane, pas - post-acrosomal sheath

**Fig. S2.**
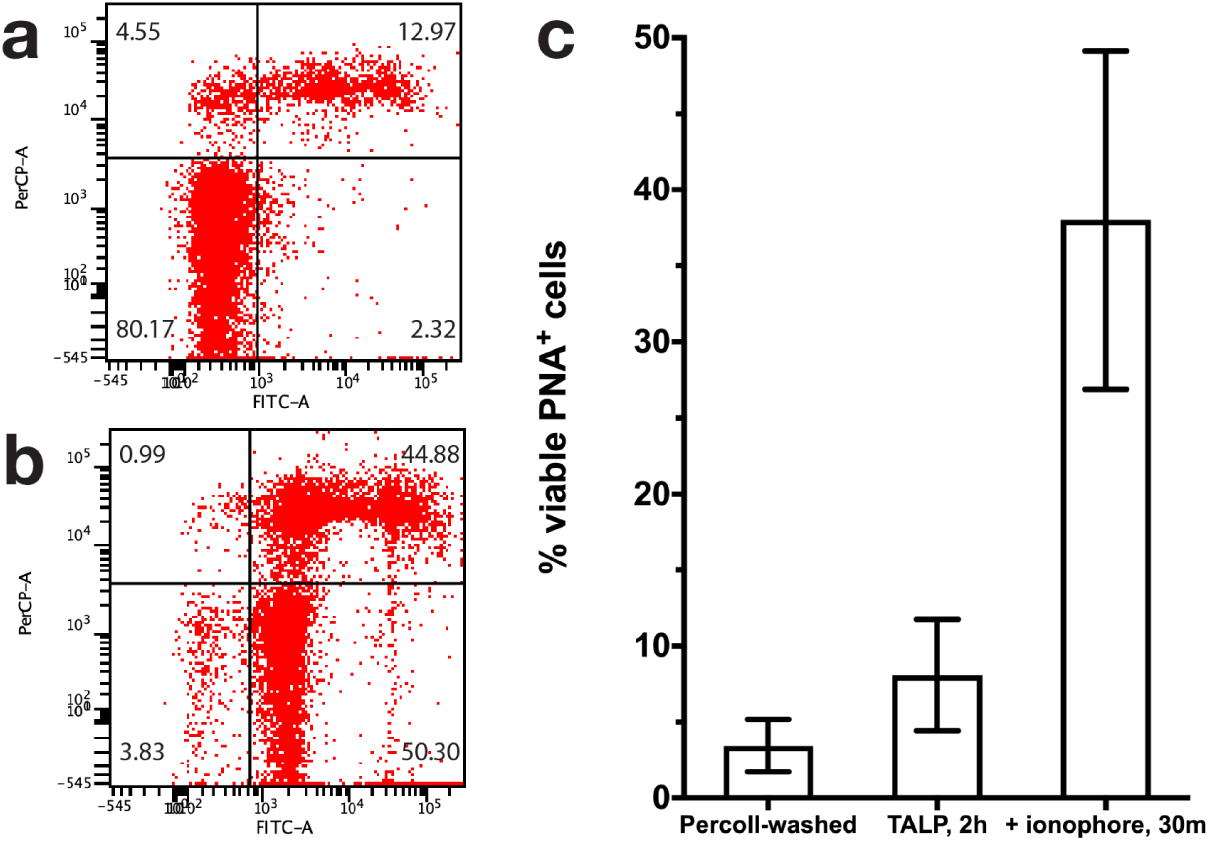
Flow-cytometric assessment of acrosome reaction efficiency. **(a-b)** Flow cytograms of non-capacitated sperm (a) and sperm stimulated to acrosome react with ionophore A23187 (b). **(c)** Proportion of viable acrosome-reacted sperm (PI-, PNA-FITC+) as assessed by flow cytometry. Bar graphs show mean ± standard deviation and summarize three independent experiments on three separate animals.

**Fig. S3.**
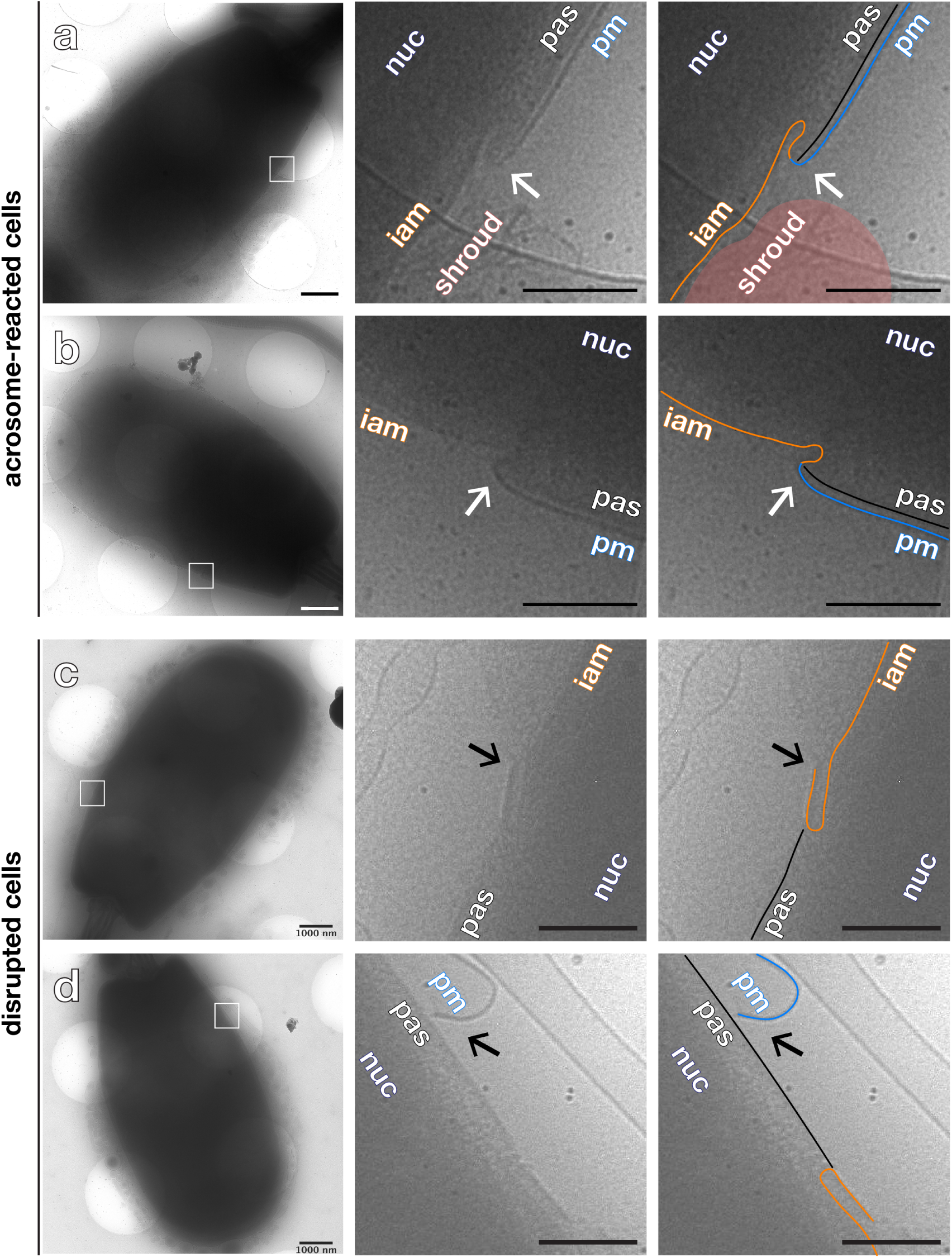
Distinguishing acrosome-reacted sperm from membrane-disrupted sperm in low-magnification cryo-EM projection images. Low-magnification high-dose cryo-EM projection images of acrosome-reacted **(a-b)** and membrane-disrupted **(c-d)** sperm. Note how acrosome-reacted sperm successfully re-seal at the equatorial/post-acrosomal region (a-b, white arrows in digital zooms); in contrast, membrane-disrupted sperm have also lost the plasma membrane overlying the post-acrosomal sheath (c-d, black arrows in digital zooms). Also note how the apical region becomes very thin in acrosome-reacted cells; in contrast, membrane-disrupted sperm retain an acrosomal ghost around their heads. **Scale bars:** (a-d) 1 µm; digital zooms: 250 nm. **Labels:** pm – plasma membrane, oam – outer acrosomal membrane, iam – inner acrosomal membrane, pas – post-acrosomal sheath, nuc – nucleus, shroud – acrosomal shroud. **Color scheme:** blue – plasma membrane, orange – inner acrosomal membrane, black – post-acrosomal sheath, red – acrosomal shroud.

**Fig. S4.**
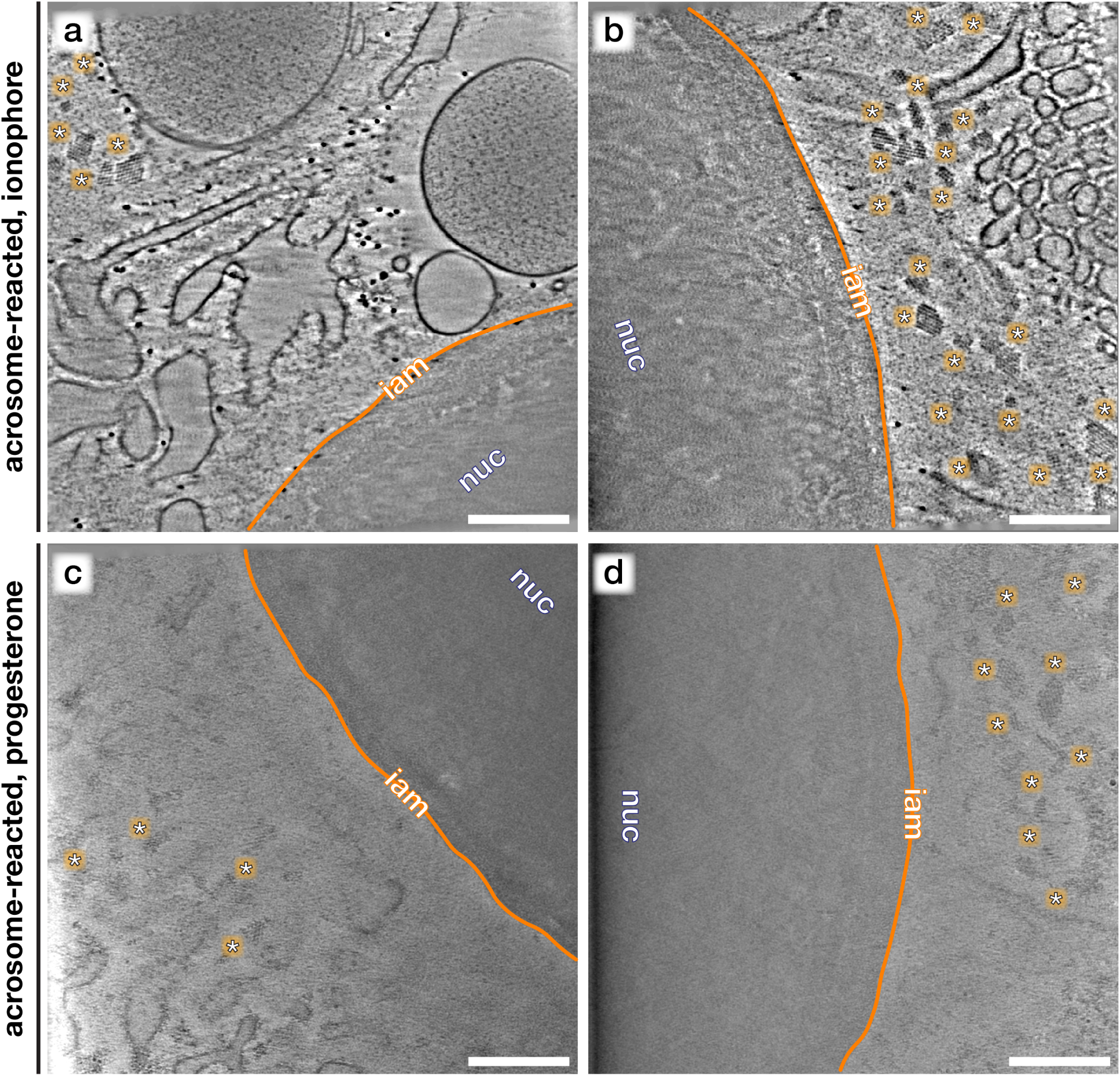
Additional examples of crystalloid patches in the acrosomal shroud of acrosome-reacted cells. Computational slices through Volta phase plate cryo-tomograms of the acrosomal shroud collected at the apical region of sperm stimulated to undergo acrosomal exocytosis with either calcium ionophore **(a**,**b)** or progesterone **(c**,**d)**. Note the presence of crystalloid patches (asterisks) of varying shapes and sizes in all tomograms. **Scale bars:** 250 nm. **Labels:** iam - inner acrosomal membrane, nuc - nucleus.

**Fig. S5.**
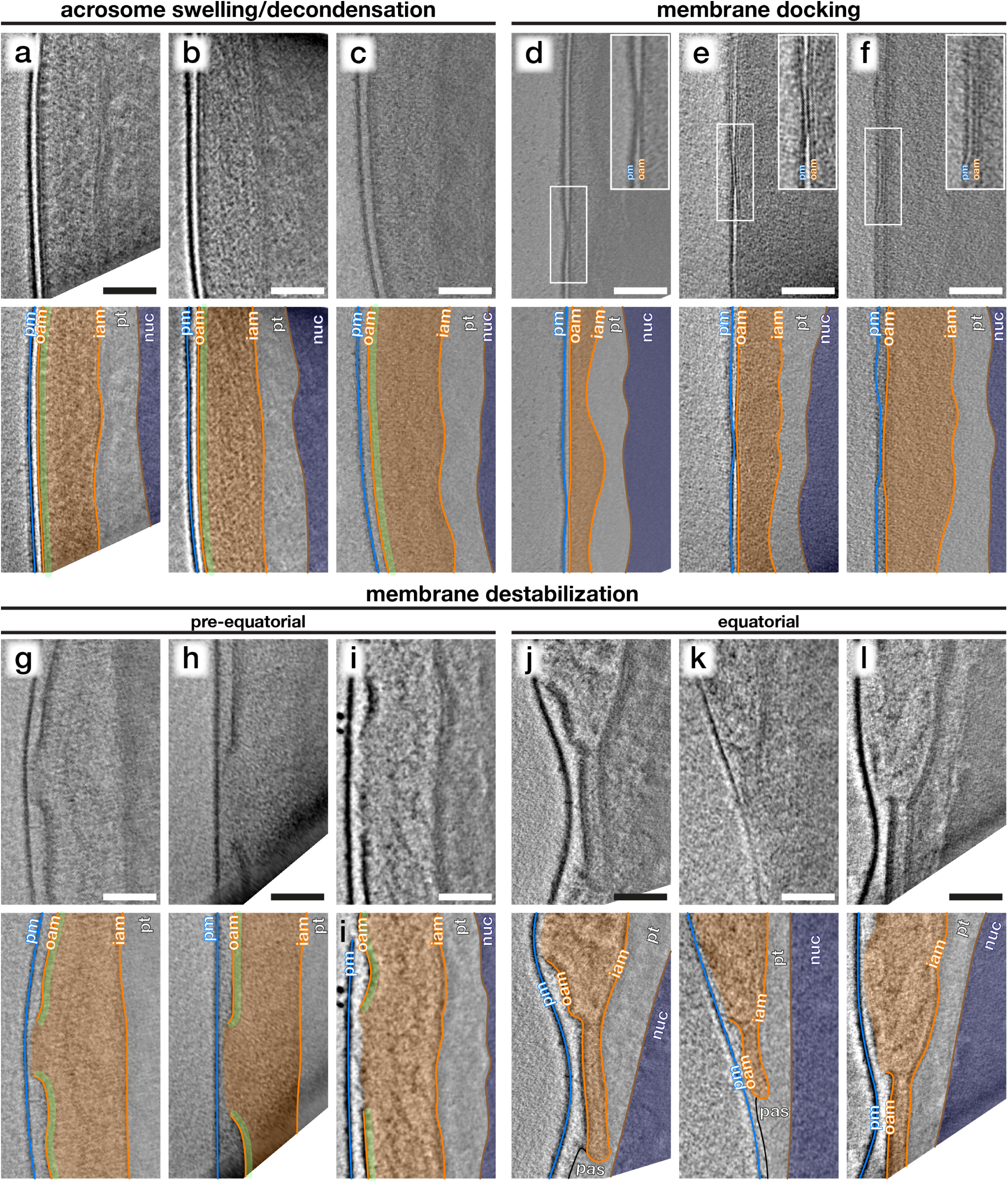
Additional examples of membrane remodelling intermediates observed in sperm incubated in capacitating media without ionophore stimulation. Computational slices through cryo-tomograms of sperm with **(a-c)** swollen acrosomes and intact membranes; **(d-f)** swollen acrosomes and locally-docked membranes (insets); and **(g-l)** profusely-swollen acrosomes and destabilized membranes at the pre-equatorial (g-i) and equatorial regions (j-l). Tomograms in (a), (b), (d), (i), (j), and (l) were acquired with the Volta phase plate while tomograms in (c), (e), (f), (g), (h), and (k) were acquired with defocus-contrast. **Scale bars:** 100 nm. **Labels:** pm – plasma membrane, oam – outer acrosomal membrane, iam – inner acrosomal membrane, pt – perinuclear theca, pas – post-acrosomal sheath, nuc – nucleus. **Color scheme:** blue – plasma membrane, orange – outer and inner acrosomal membranes, green – membrane protein densities, grey – perinuclear theca and post-acrosomal sheath, brown – nuclear envelope, dark blue - nucleus.

**Fig. S6.**
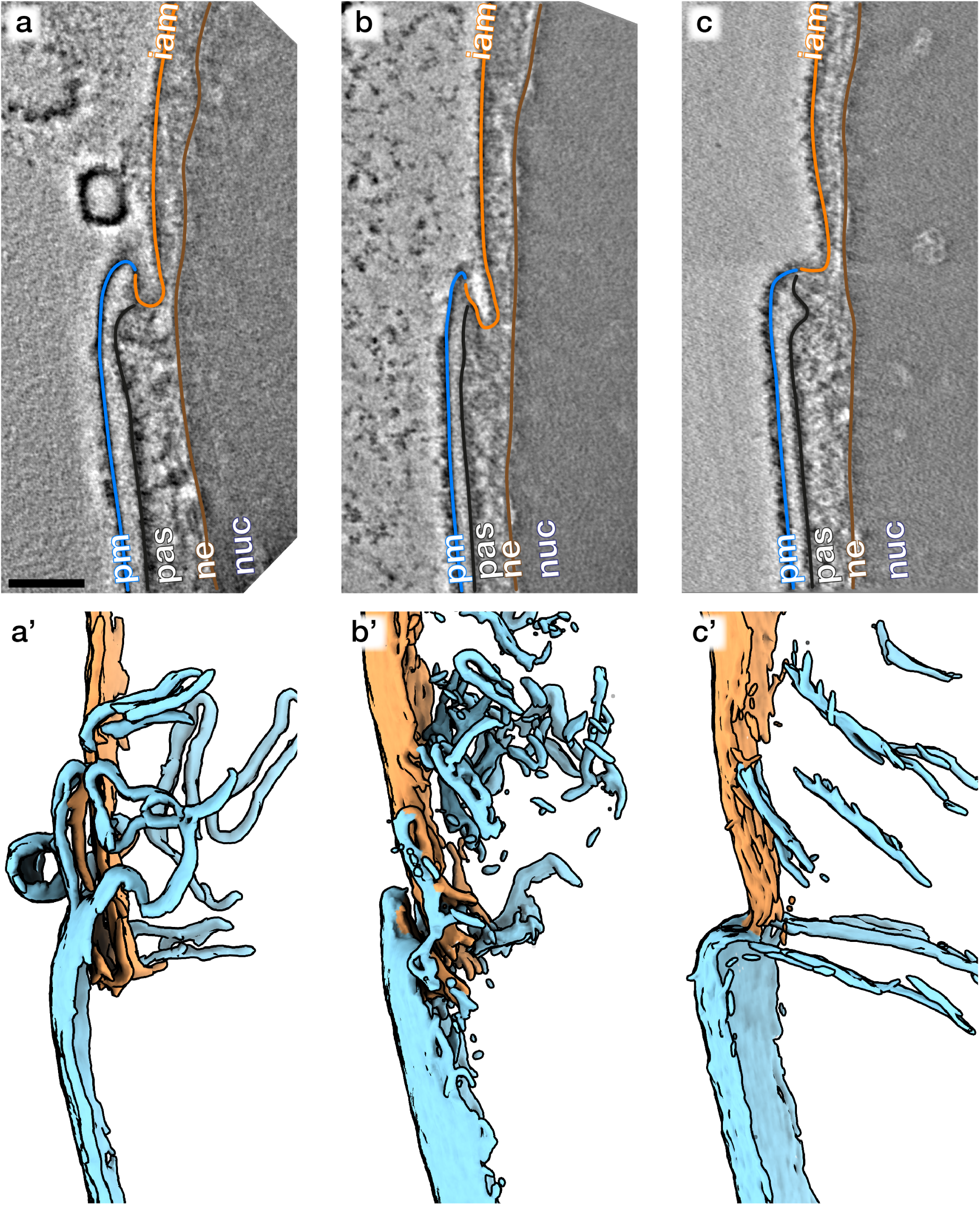
Additional examples of the remodelled topography of acrosome-reacted cells. **(a-c)** Computational slices and **(a’-c’)** corresponding three-dimensional segmentations of Volta phase plate cryo-tomograms of acrosome-reacted sperm. **Scale bars:** 250 nm. **Labels:** pm – plasma membrane, iam – inner acrosomal membrane, pas – post-acrosomal sheath, ne - nuclear envelope, nuc – nucleus. **Color scheme:** blue – plasma membrane, orange – inner acrosomal membrane, black – post-acrosomal sheath, brown – nuclear envelope

**Fig. S7.**
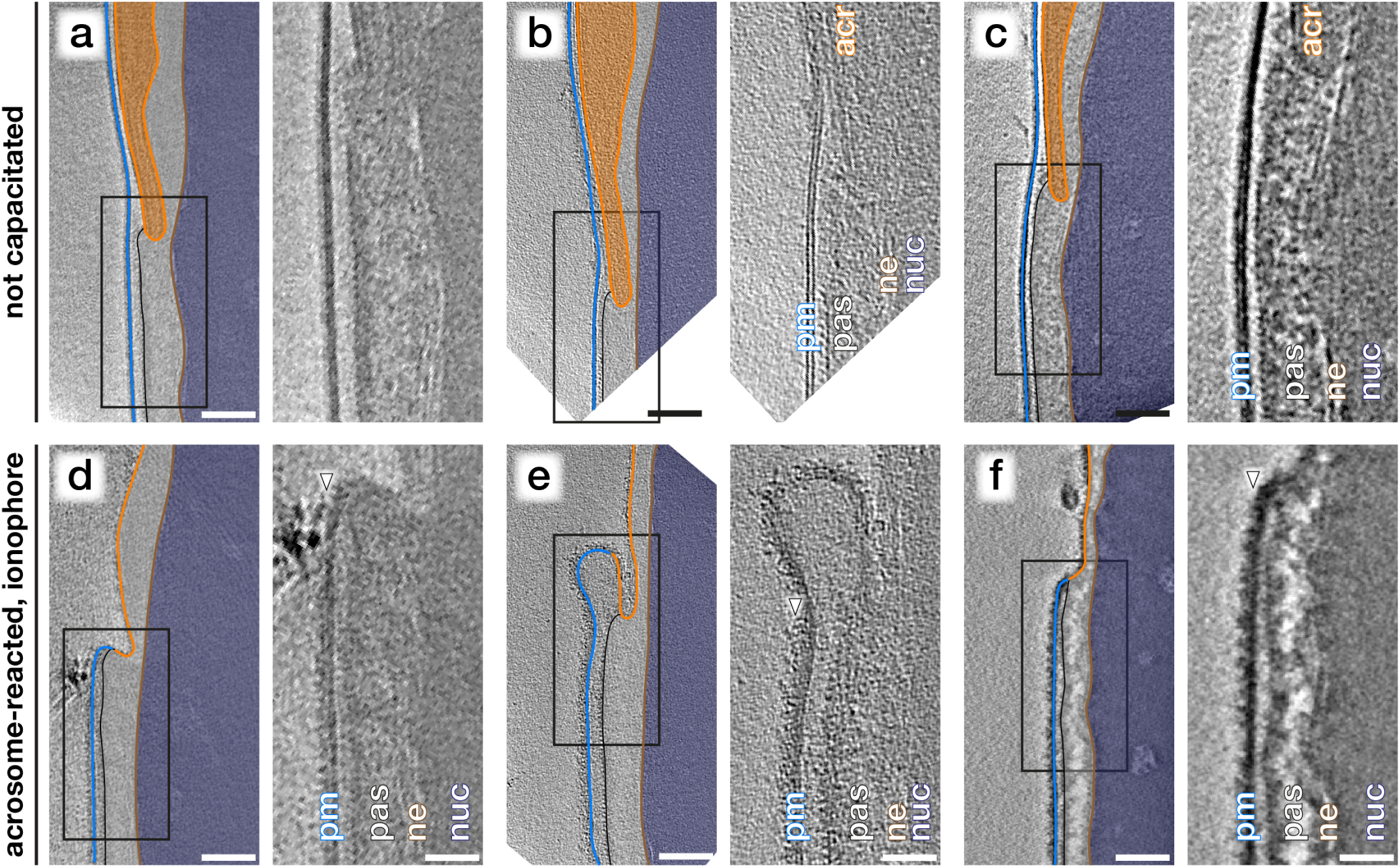
Additional examples of membrane protein relocalization onto the post-acrosomal plasma membrane of acrosome reacted cells. Computational slices through cryo-tomograms of the equatorial/post-acrosomal region in unstimulated, non-capacitated **(a-c)** and acrosome-reacted **(d-f)** sperm. The change in membrane protein decoration (arrowheads in d-f) is seen consistently even in three different imaging conditions: intermediate-magnification defocus contrast (a,d), high-magnification defocus contrast (b,e), and high-magnification Volta phase plate contrast (c,f). **Scale bars:** 100 nm. **Labels:** pm – plasma membrane, iam – inner acrosomal membrane, pas – post-acrosomal sheath, ne - nuclear envelope, nuc – nucleus. **Color scheme:** blue – plasma membrane, orange – inner acrosomal membrane, black – post-acrosomal sheath, brown – nuclear envelope

**Table S1.** Dataset summary reporting number of tomograms in the dataset representing each membrane remodelling inter-mediate.

| description | condition |  |  |
| --- | --- | --- | --- |
|  | unstimulated | capacitated | acrosome-reacted |
| dense acrosome, membranes intact | 26 | 10 | - |
| swollen acrosome, membranes intact | - | 15 | - |
| swollen acrosome, membranes destabilized | - | 16 | - |
| acrosome-reacted, apical region, shroud present | - | - | 13 (ionophore), 5 (progesterone) |
| acrosome-reacted, apical region, shroud absent | - | - | 6 (ionophore) |
| acrosome-reacted, equatorial region | - | - | 37 (ionophore) |
| **total number of tomograms** | **26** | **41** | **56** (ionophore), **5** (progesterone) |
| **total number of animals** | **7** | **5** | **6** (ionophore), **1** (progesterone) |

